# A Convex Point Process Model of Heartbeat Dynamics for Inference, Prediction, and Information Quantification

**DOI:** 10.1101/2025.08.28.672903

**Authors:** Andrew S. Perley, Matthew Martinez, Tommaso Mercadante, Sabrina Liu, Todd P. Coleman

## Abstract

The dynamics of heartbeat intervals provide important insights into cardiovascular and autonomic nervous system function. Conventional analytical approaches often use fixed-window averaging, which can obscure rapid changes and reduce temporal resolution. Point process models address this limitation by operating in continuous time, enabling more precise characterization of heartbeat variability. A landmark example is the history-dependent inverse Gaussian (IG) point process model of Barbieri et al. (2005), which captures temporal dependencies in heartbeat timing. However, the nonconvex likelihood of the IG model complicates parameter estimation, requiring careful initialization and adding computational burden. In this work, we introduce a convex alternative: a history-dependent gamma generalized linear model (GLM) for heartbeat dynamics. Applied to a tilt-table dataset, our approach yields accurate and robust heart rate estimation. We further extend the model to two more applications: (1) sequential prediction of interbeat intervals, outperforming common machine learning algorithms, and (2) computation of information-theoretic measures demonstrating its utility in quantifying the influence of cardiac medications on heartbeat dynamics.

## Introduction

Understanding heartbeat dynamics has been central to the study of the autonomic nervous system (ANS), particularly in how it modulates physiological responses related to sleep, emotion, pain, and stress in both health and disease contexts (1). The ANS regulates involuntary physiological functions and interacts closely with the cardiovascular system (2). Since beat-to-beat fluctuations in heart rate reflect the dynamic balance of sympathetic (“fight-or-flight”) and parasympathetic (“rest-and-digest”) activity, heart rate variability (HRV) serves as a noninvasive window into real-time autonomic function (2).

Despite the rich physiological information embedded in heartbeat intervals, conventional HRV analysis methods often obscure fine temporal details. Many rely on windowed averaging or spectral decomposition, which limits their ability to resolve second-to-second changes in autonomic tone (3). In contrast, point process models preserve the temporal structure of cardiac activity by modeling heartbeats as discrete events occurring in continuous time, aligning more closely with the electrophysiological mechanisms of cardiac pacemaker cells (1).

A seminal contribution to this line of work is the point process framework proposed by Barbieri et al. (4), which models the evolution of the cardiac membrane potential as a Gaussian random walk with drift, leading to an inverse Gaussian distribution for interbeat intervals (RR intervals). While this approach offers high temporal precision and physiological interpretability, its implementation poses computational challenges due to the nonconvex nature of the maximum likelihood estimation problem. As a result, model fitting is sensitive to initialization and often computationally intensive.

In this paper, we propose an alternative framework for modeling heartbeat dynamics using a history-dependent gamma generalized linear model (GLM). This formulation retains the statistical fidelity of the Barbieri model while simplifying parameter estimation by making the likelihood optimization problem convex. By reducing the dependence on initial conditions and improving computational efficiency, our approach facilitates broader adoption of point process models for analyzing autonomic function from heartbeat data.

Beyond model fitting efficiency, we expand the scope of prior work in two important directions. First, we adapt the gamma-GLM framework to a sequential prediction setting, enabling real-time forecasting of upcoming interbeat intervals. Second, we demonstrate how this model can be used to compute information-theoretic measures between heartbeat dynamics and external covariates—in particular, pharmacological interventions. Using data from an experiment testing the effects of varying cardiac drugs, we show that our method can differentiate between the effects of distinct cardiac medications based solely on observed cardiac state, providing a potential tool for evaluating drug responses in vivo.

The remainder of this paper is structured as follows. We begin with a brief Preliminaries section to introduce notation and some information-theoretic quantities. In the Methods section, we describe the construction of the gamma-GLM model and the validation procedures used to compare it against the inverse Gaussian framework. In Results, we evaluate model performance in terms of goodness-of-fit, computational efficiency, sequential prediction accuracy, and information-based drug classification. Finally, in the Discussion, we examine the broader implications of our approach and outline possible directions for future research.

### A. Preliminaries

#### A.1. Notation

For any random variable, let the uppercase (e.g. *Y*) represent the random variable itself, and the lowercase (e.g. *y*) represent a realization of that random variable. Vectors are denoted with an underline (e.g. *x*), matrices in bold (e.g. **A**), and scalars are left just as the variable name.

#### A.2. Definitions

*Definition 1* (Kullback-Liebler (KL) Divergence) The Kullback-Liebler divergence, also known as the relative entropy, from *P* to *Q* is defined as

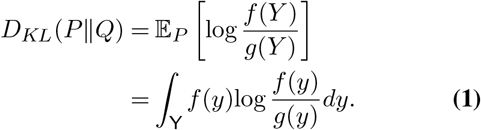

Although relative entropy is not a true distance metric—since it lacks symmetry—it is commonly used as a measure of how different two probability distributions are. It can also be interpreted as the expected penalty incurred when encoding data under distribution *Q* while the data is actually generated according to distribution *P* (5).

Now consider a pair of random variables, *X* and *Y*, with joint distribution *P*_*X,Y*_ and marginal distributions *P*_*X*_ and *P*_*Y*_, respectively.

*Definition 2* (Mutual Information) Let *X* and *Y* be random variables with joint density *f*_*X,Y*_ (*x, y*). The mutual information between *X* and *Y* is defined as:

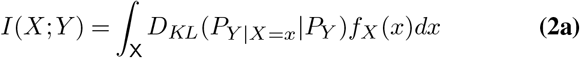

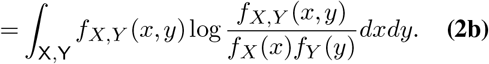

Mutual information is a symmetric quantity and often interpreted as the amount of information one random variable provides about the other (5).

## Methods

### B. A History-Dependent Gamma GLM Point Process Model

Prior point process heartbeat dynamics work utilizes a history-dependent inverse-Gaussian (IG) distribution, motivated by how an IG models the first-passage times of a random walk (4). However, this framework results in a nonconvex fitting procedure without guarantees of a model fitting procedure obtaining the maximum likelihood estimate. In this section, we will propose to use the gamma distribution as a natural analogue to recast the problem using a convex formulation. Throughout this paper, we will consider the problem of modeling the timings between heartbeats. Since the time between heartbeats is nonnegative, we must choose an appropriate distribution to accommodate the nature of these timings. Here we will choose to model our continuous nonnegative random variable *Y* with a gamma distribution with shape and rate parameters (*α, β*) as:

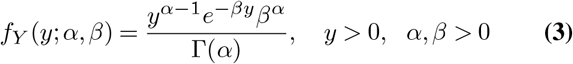

where Γ(·) is the gamma function. The above model is useful in modelling the variation in single heartbeat timings, however, we are interested in understanding the statistical structure in a time series of heartbeat timings.

Define the random vector, *Y* ≜ (*Y*_1_, *Y*_2_, …, *Y*_*n*_), as the interbeat intervals (in seconds) between successive heartbeats, where any *Y*_*j*_ ∈ ℝ_+_ is nonnegative. Here we will use the timings between R peaks in an electrocardiogram, i.e., the RR interval, as our interbeat interval. We know from physiology that an RR interval is statistically dependent on recent past sympathetic and parasympathetic inputs to the heart (4). As such, we model each RR interval *Y*_*j*_ as statistically dependent on its recent history ℋ _*j*_ = (*Y*_*j*−1_, …, *Y*_*j*−*p*_). A natural way to do this is by constructing a gamma GLM as follows:

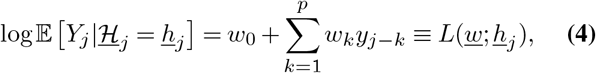

where *w*_0_, *w*_1_, …, *w*_*p*_ are a fixed set of weights that connect the expectation of *Y*_*j*_ to its history of *p* lags into the past. The log link is used to map 𝔼 [*Y*_*j*_ | ℋ _*j*_ = *h*_*j*_ ∈ ℝ_+_ ] onto ℝ, since *w*_0_, *w*_1_, …, *w*_*p*_ can take on any real value. This then describes a point process model where the distribution of any RR interval conditioned upon its past is described by a gamma distribution.

### C. Model Fitting

For the gamma density with parameters (*α, β*), we have from Eq. (3) that

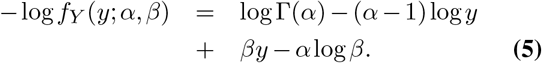

Given the expectation of a gamma random variable *Y* is 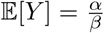, we have that for a gamma GLM with log link function:

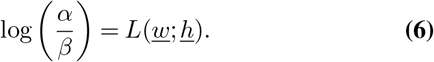

We note from the above equation that the model can alternatively be parameterized by *α* and *w*.

We first consider the case where there is one RR interval of duration *y* and history *h*. Then the negative log likelihood described in terms of shape parameter *α* and weights *w* is given by

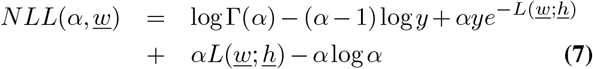

Then for any *α >* 0, the maximum likelihood (ML) estimator for *w* is given by

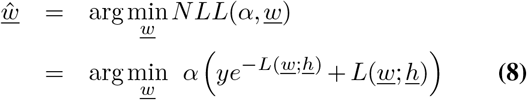

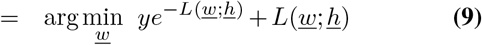

where Eq. (8) follows from elimination of terms in the sum that do not depend on *w* and Eq. (9) follows from the fact that *α >* 0.

If we now have *J* heartbeat intervals, the ML estimator for *w* is given by

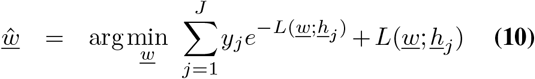

Note that this minimization problem is convex in *w* and does not force weights to be nonnegative. It thus overcomes two limitations of the IG approach (4).

After finding the ML estimate for *w*, we can find the ML estimate for *α* as follows. Define *L*_*j*_ ≜ *L*(*ŵ*; *h*_*j*_). Taking into consideration all terms from the NLL that involve *α*, we have from that:

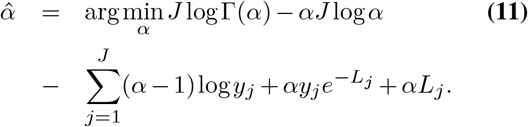

The problem described in Eq. (12) is also convex, thus making the full model fitting procedure a two-step procedure where both steps are separate convex optimization problems. Since we have both 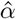 and *ŵ*, we can directly compute 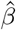 through Eq. (6).

### D. Goodness of Fit

To assess goodness of fit of our model, we follow the time-rescaling methodology first proposed by Brown et al., for neural spiking dynamics and adapted by Barbieri et al., for heartbeat dynamics through the time-rescaling theorem (4, 7). First, define the times at which R-wave events occur, 0 *< u*_1_ *< u*_2_ *<* … *< u*_*k*_ *< T*, where *T* is the total time of recording. We can express any *u*_*k*_ as the sum of all interbeat intervals leading up to *u*_*k*_,

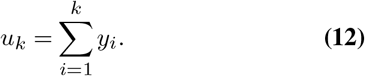

The conditional intensity of the point process can be described as (4):

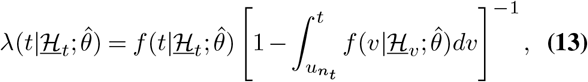

Where 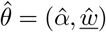, the parameters fit by the model, 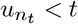, is the time at which the most recent previous R-wave event occurred, and 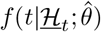 is the gamma GLM density with history ℋ _*t*_ evaluated at time 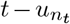, which by definition of 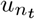 is nonnegative.

We can then compute the time-rescaled interbeat intervals by

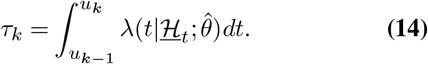

The time-rescaling theorem states that for an arbitrary point process, the time-rescaled inter-event intervals (*τ*_*k*_ : *k* ≥ 1) as defined above are independent exponentially distributed random variables with unit rate (7). We can assess the goodness of fit by the transformation *Z*_*k*_ = *F*_exp_(*τ*_*k*_), where *F*_exp_(*s*) = 1 − *e*^−*s*^ is the cumulative distribution function (CDF) of a unit rate exponential random variable. Since it is wellknown that inputting a random variable into its own CDF produces a uniform random variable on [0, 1], (*Z*_*k*_ : *k* ≥ 1) should be uniformly distributed on [0, 1] if the data was distributed according to the proposed model 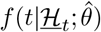. We can compare (*Z*_*k*_ : *k* ≥ 1) to the uniform random variable on [0, 1] by means of a Kolmogorov-Smirnov (KS) plot. We then calculate the KS distance between the empirical CDF of (*Z*_*k*_ : *k* = 1, …, *n*) and compare it to the theoretical CDF of the uniform distribution *F*_*U*_ (·) as:

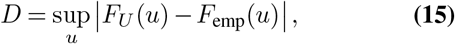

where *F*_*U*_ (·) is the CDF of a uniform random variable on [0, 1] and *F*_emp_(·) is the empirical CDF of the *z*_*k*_. This metric measures the level of dissimilarity between the distributions.

### E. Sequential Prediction of Subsequent Heartbeats

One interesting application of statistical models is the ability to perform prediction, to use past information to predict future occurrences. One way to perform such a task is via sequential prediction, which uses principles of information theory and game theory to consider algorithms that are robust to worst case scenarios (8). In this paper we will utilize a form of sequential prediction where we attempt to construct a probability distribution that will characterize a future data point based upon past datapoints. We choose this form because of theoretical guarantees the worst case performance, or regret, with respect to a class of possible prediction algorithms. The specific approach we take can be described as a sequential solution to optimization problems that leverages how the class of models we use come from the exponential family (9).This form of sequential prediction problem can be described as the following optimization problem:

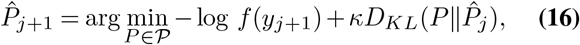

where *f* is the probability density function associated with probability distribution *P*, 𝒫 is the set of possible probability distributions to which *P* can belong, 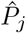 is the predictive probability for interval 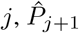 is the inferred predictive distribution of heartbeat interval *j* + 1, and *κ* is a regularization parameter that helps constrain how much the new distribution 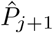 can deviate from 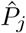 in trying to predict the next interval (9). In this problem, we adapt this method to our specific history-dependent gamma GLM and let 𝒫 be the set of all gamma GLMs with history length *p*. As such, we can rewrite the problem as an optimization problem over the parameters of the GLM:

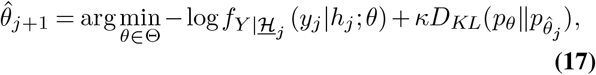

where *p*_*θ*_ has density 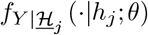 and Θ is the set of all possible *θ*_*j*_. Since we use a gamma GLM, the conditional distributions specified by fit models are gamma distributions. As such, for two instances of parameters, they induce two distinct gamma distributions *P*_1_, with parameters (*α*_1_, *β*_1_) and *P*_2_, with parameters (*α*_2_, *β*_2_). Since both are gamma distributions, and are thus members of the exponential family, a closed form solution for the KL divergence between two gamma distributed *P*_1_ and *P*_2_ exists and can be written in terms of the parameters as follows (10):

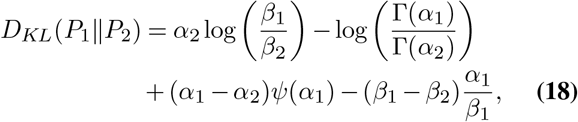

where Γ(·) is the gamma function and *ψ*(·) is the digamma function. Since the *β*s are a function of the model weights and *α*, we can recast this in terms of the model weights and the *α*s using Eq. (6).

### F. Extension to Information Measures

Gamma GLMs can be expanded with additional regressors, such as respiratory effort or blood concentration of a drug, beyond our original history-dependent RR interval construction. In this case, we may be interested in how this additional information aids in predicting future heartbeat timings. One way to quantify the improvement in predicting future outcomes with additional side information is using conditional mutual information, which in many contexts uniquely characterizes this improvement under a natural assumption regarding data processing (11). Specifically, here we use the conditional mutual information (CMI) between the current heartbeat and the additional regressors, given the history of previous heartbeats. For the rest of this paper, we will refer to *X* ∈ ℝ^*R*^ as the random vector of *R* additional regressors. Thus, we can compute the CMI by the following equation:

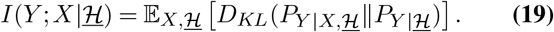

One advantage of this representation is that if we represent the density of *P*_*Y |X*, ℋ_ as a gamma GLM, we can compute the KL divergence between the model with and without the additional regressors in closed form by Eq. (18). This is because the gamma GLM link equation in Eq. (4) associated with *P*_*Y* |*X*, ℋ_ can be expressed as

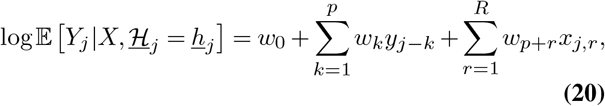

where the *w*_*p*+*r*_ represent the additional regression weights and *x*_*j,r*_ represents the *r*^*th*^ component of *X*_*j*_. Given this representation, we can solve Eq. (10) and Eq. (12) and estimate 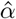 and *ŵ* as described earlier. We can then transform this into a 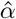 and 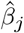 in accordance with Eq. (32). Thus, since Eq. (18) relies only upon the *α* and *β* values of each distribution, we can compute the KL in closed form between the models with and without additional regressors. On the other hand, computing the outer expectation is, in general, not closed form. As such, here we approximate the outer expectation using a sample average over the data distribution. That is,

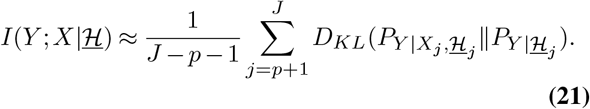

Borrowing from causality principles in economics, one can interpret this measure as “how much better can I predict a future heartbeat timing knowing additional information, if I already had access to the past heartbeat timings?” (12). That is, if we already have access to a subject’s heart rate history, can we perform even better prediction through measuring other covariates? When the CMI is large, it implies that we can gain significant information about the heartbeat dynamics from other regressors. If the CMI is 0, then it implies that, given the heart rate history, the current heartbeat and the covariates are conditionally independent. Since we are taking this expectation over the data distribution, we will refer to this method, less formally, as the KL Divergence analysis in the rest of this paper.

### G. Constructing a Predictive Model

Our GLM-based model of heartbeat dynamics enables predictive modeling via decision theory, supporting applications such as physiological state classification and real-time cardiovascular state estimation using regression (13). For example, one may be interested in predicting the emotional state of a subject using classification or estimating respiratory effort using regression (14, 15). Here, we demonstrate formulations for both classification (discrete-state prediction) and regression (continuous-state prediction) using our GLM formulation, but will only demonstrate classification in experiment later in the paper.

To formulate the problem, we first consider a generic loss function *l*(*d, x*) that determines the inadequacy associated with a decision *d* ∈ 𝒟 if the true variable of interest is *x* ∈ 𝒳. We here consider settings where we cannot measure *X* but can measure *Y* and side information *H* that both are correlated with *X* through a joint distribution *P*_*X,Y*, ℋ_, meaning that *d* must only be a function of *Y* and ℋ, namely *d* = *π*(*Y*, ℋ) for some function *π*. The quantity to minimize is given as

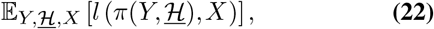

The optimal decision, attaining the Bayes risk, can be taken by minimizing the conditional expectation (16, 17):

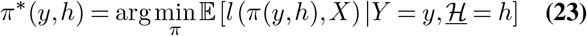

In the classification setting, for 𝒟 = 𝒳 = {1, …, *K*} pertaining to *k* classes, we consider the zero-one loss function given by *l*(*d, x*) = 1_*d/*=*x*_, where we incur a loss of 1 when the class prediction is not the same as the true class label. In this setting it can be shown that attaining the Bayes risk pertains to employing Bayes rule:

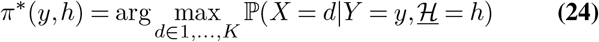

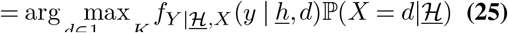

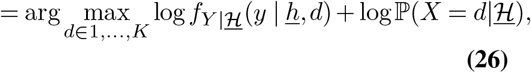

where Eq. (25) follows from Bayes’ rule (17). The corresponding optimal class is then given by the decision rule as

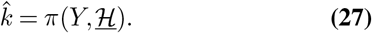

In practice, to train the classifier, we partition the training data according to class labels and fit a separate GLM to each subset. This yields class-specific parameter estimates 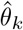, which characterize the conditional dynamics of heartbeat features under each physiological state. We assume in this paper that the prior probability ℙ (*X* = *k* | ℋ) is uniform over all classes, which gives us the maximum likelihood decision rule. Thus, given a new observed heartbeat timing and history (*Y*, ℋ), we compute its likelihood under each class model *f*_*Y*_ | ℋ (*y* | *h, k*), and assign the class that maximizes this likelihood according to Eq. (27). For a finite dataset of timings and histories, {(*Y*_1_, ℋ _1_), …, (*Y*_*N*_, ℋ_*N*_)}, we can assign the optimal class using the sample average of the log likelihood,

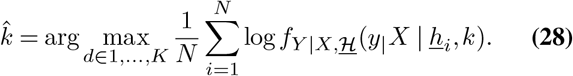

In the continuous setting for posterior regression, we instead consider the squared loss function on the additional regressors, *X*, as 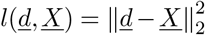, where we desire our decision rule *π*(·) to operate on *Y* and ℋ to output some prediction in the space of 𝒳. This can be computed from the posterior distribution on *X* given by Bayes’ rule as

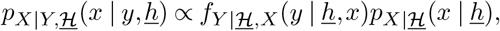

where we will need to assume some form of the prior distribution *p*_*X* |ℋ_ (*x* |*h*). Since posterior inference can be computationally intensive, we will not demonstrate an example of this here. However, we still show this formulation as a potential future application of our model for practitioners.

## Experiments

### H. Assessment of Performance

To assess the performance of the proposed algorithm compared to the existing algorithm, we collected a healthy human dataset of over 7000 RR intervals from and fit both algorithms to the data. The data was collected from a human subject wearing an ambulatory electrophysiology monitoring device as described in Gharibans et al (18). Both the amount of time it took to fit the algorithm and the KS distance were calculated to assess computational time and model fit. Both lower runtime and lower KS distance indicate a better performing algorithm. Since the initial portion of a recording can tend to have unreliable measurements, we chose a window of 120 seconds at the beginning to exclude from the analysis. We chose a model order, *p* = 6, as in Barbieri et al., by the Akaike Information Criterion (AIC) for the IG model, and set it to be the same for the gamma model for comparison (4, 19). We then ran a sweep to fit models for *J* ∈ [100, 7250]. Specifically we ran the model for *J* = 100 and increased 100 intervals up to *J* = 1000, then every 250 intervals up to *J* = 7250.

### I. Heartbeat Time Series Inference

As compared to traditional techniques to estimate heart rate by performing 60*/RR* for each RR interval, point process heartbeat models have an ability to infer more robust ‘instantaneous’ heart rate (HR) time series. Here, we use the mean HR as computed by the conditional expectation of the model given its past history. In particular, since our model is over the RR intervals, we adapt it to heart rate, by deriving the distribution of the random variable *HR* = 60*/Y*. This is characterized by an inverse gamma distribution with mean

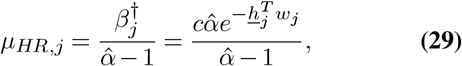

where *c* = 60 s*/*min converts the mean from beats per second to beats per minute. Note that for every history *h*_*j*_, there exists a conditional mean heart rate, thus a mean heart rate can be inferred on a beat-by-beat basis for which we have a history. In this experiment, we use a tilt-table dataset, where subjects undergo an experiment where they are laid flat and tilted vertically at varying rates (20–22). This experiment is commonly used to assess the autonomic function of participants. We choose this dataset to demonstrate the ability of our model to track heart rate in fast changing physiological states. Here we use *p* = 2 as a fixed history length for all subjects in the tilt table dataset. For more judicious choice of history length per subject, we could choose to use AIC (19). However, here we show that even with a small history this model produces more robust estimates of the heart rate time series.

### J. Sequential Prediction vs. Machine Learning methods

In this experiment, we examine the ability of our model to do next-beat heartbeat timing predictions given prior data. We compare against three standard machine learning methods: random forest regression, support vector regression (SVR), and multilayer perceptron (MLP) regression, all implemented using the scikit-learn python package (23). We specifically choose these methods as comparisons, since our datasets are small and the proposed strength of our model is that we can do inference in a subject-specific manner as shown in prior literature (24). The random forest regression model was fit using 100 decision trees with a max depth of 3. The support vector regression was fit using a polynomial kernel, a regularization constant, *C* = 1, and *E* = 0.1. The MLP regression model was set with 3 hidden layers each with 10 ReLu units, an adaptive learning rate starting at 0.01, an L2 regularization parameter of 0.01, and was optimized using the ADAM optimizer. All other parameters in these algorithms were left as the default settings in the scikit-learn package. For the machine learning methods, we use the history data as the input and the next proceeding RR interval as the output. We use 300 RR intervals, approximately 5 minutes, of data as training data for each model and the remainder of the recordings as test data. For all models, we fit histories for *p* = 12, 24,…, up to half the length of the training data and choose the fit with the lowest training MSE for each model class. We use the same tilt table dataset from the last experiment to assess both the mean absolute error (MAE) and mean square error (MSE) on test data in prediction performance. This dataset offers the ability to understand prediction errors when the heartbeat timings are locally stationary and experience abrupt changes in statistics.

### K. Influence of Cardiac Drugs on the Heart

We evaluated our model’s capacity to capture the effects of various drugs, presenting a possible clinical application of this algorithm. The data collected for this assessment was a randomized, double-blind, clinical trial of 22 healthy subjects where 16 rounds of EKG results were collected in triplicate over 10-second intervals as described in Johannesen et al (22, 25). The 16 rounds were collected as follows for each drug: 1 point pre-dose (-0.5 h) and 15 points post-dose (0.5, 1, 1.5, 2, 2.5, 3, 3.5, 4, 5, 6, 7, 8, 12, 14, and 24 h). The drugs assessed in this trial were Dofetilide (500 µg), Quinidine (400 mg), Ranolazine (1500 mg), and Verapamil (120 mg) and a placebo was included for the fifth period of data collection. All of these drugs are QT prolonging drugs, so our model’s ability to distinguish between each drug would demonstrate its high sensitivity when comparing drugs that are reported to have the same clinical effect.

To assess the amount of additional information provided by the drug identity, we compute the CMI as described in Eq. (21). To compute the CMI, we need to be able to compute the KL divergence at each beat time between a model that includes the additional regressor and one that does not include it. Thus, in the following section we will describe our method of constructing both models and computing the appropriate KL divergences.

We begin by preprocessing the heartbeat information, isolating the RR intervals from the EKG data collected from Johannesen et al (25). QRS complex peaks are detected using a Teager Kaiser Energy Operator (TKEO) based method described in Beyramienanlou et al. (26). RR intervals were then computed as the difference in timings between successive QRS complex peaks.

Then for each subject, we construct two GLM point process models with and without the drug. For each of these models, since each subject has 3 EKGs at 16 time points for each drug, we need to take care in how we pool all this data to fit a single model with drug information and a single model without drug information. To do so, we will represent the problem equivalently in terms of the corresponding regressor matrix. Consider Eq. (4), and let

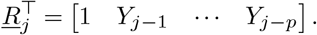

Then Eq. (4) can be equivalently represented as the inner product

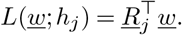

Given this, we can then rewrite Eq. (10) as

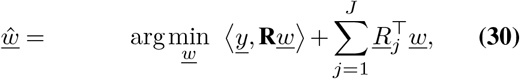

The model parameters 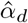 and *ŵ*_*d*_ are solved by the same Eq. (30) and Eq. (12). For each of the models, with and without the explicitly encoded drug information, we denote their regressor matrices as 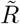 and *R* respectively. The specific construction of these regressor matrices is left in Appendix A. To compute the KL divergences needed for Eq. (21), we can then take advantage of Eq. (6), where we can, for each row of the full regressor matrix 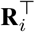, compute a *β*_*i*_ value as

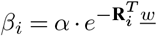

Thus for each history, as represented by a row of the full regressor matrix, we can use the global 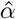 parameter and 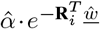, and the corresponding 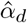 and 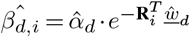, and plug into Eq. (18) to compute a KL divergence between the models for the history at row *i*. Supposing that both *R* and 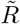 have *T* rows, then we compute the total CMI as the sample average of the KL divergences:

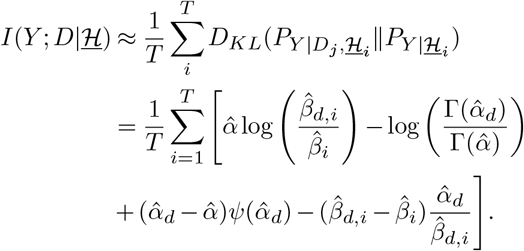

Although outside the scope of this publication, in dynamic settings where it would be desirable to track this CMI as the history changes, using sequential prediction to compute both entries in the KL divergence could be done, allowing for a dynamic measure of this information content (27). Because these values depend on the random data points, we here consider an approach to determine statistical significance. Specifically, we use a technique called permutation testing, a type of surrogate data analysis. The idea is to break the original structure in the data by randomly shuffling it, and then observe samples of CMI values obtained just by chance.

For this experiment, we randomly shuffle the rows of *R* and 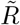, re-fit the model using these shuffled inputs, and calculate the resulting CMI to generate one sample of CMI produced by chance. We repeat this process 10,000 times to create a monte carlo estimate of the null distribution i.e., a distribution of CMI values we’d expect if there were no real relationship in the data.

The key assumption is that shuffling breaks the connection between each heartbeat and its true history or drug input, so any meaningful structure in the data is lost. If the actual (unshuffled) CMI is much larger than most of the values from this null distribution, we deem it statistically significant.

We use a significance level of 0.05 and apply a Bonferroni correction to adjust for multiple comparisons.

### L. Predicting Drug Identity from Heartbeats

We expand on our work on information measures by building a predictive model that can predict the drug taken based on differences in heartbeat intervals, history parameters, and condition parameters. To do so, we construct the model by implementing a 2:1 train-test split, where two of the three simultaneous EKG recordings are used for model training, and the last is used for the testing phase. We one-hot encode for each drug trial condition by implementing the following formula:

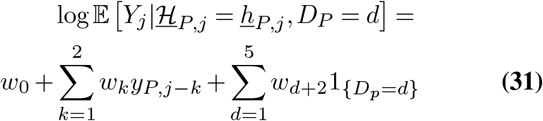

These regressor matrices were constructed based on the training data for each drug trial, and the alpha terms for each matrix were calculated using the negative log likelihood function described in Eq. (12). Five separate test regressor matrices were constructed by including the offset term, two prior intervals, and hard encoding for each condition with the arrays in table 1.

**Table 1.** Fixed one-hot encoding of each drug.

| Drug | Hard Encoding |
| --- | --- |
| Placebo | $\begin{bmatrix} 0 & 0 & 0 & 0 \end{bmatrix}$ |
| Verapamil | $\begin{bmatrix} 1 & 0 & 0 & 0 \end{bmatrix}$ |
| Dofetilide | $\begin{bmatrix} 0 & 1 & 0 & 0 \end{bmatrix}$ |
| Ranolazine | $\begin{bmatrix} 0 & 0 & 1 & 0 \end{bmatrix}$ |
| Quinidine | $\begin{bmatrix} 0 & 0 & 0 & 1 \end{bmatrix}$ |

Using the *α*s calculated for each drug trial condition in the training phase and the regressor matrices for each drug prepared in the testing condition, the test values *β* were generated as shown in Eq. 32.

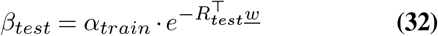

With *α*_*train*_ and *β*_*test*_ values calculated for each condition, they are applied to the heartbeat intervals reserved for the test condition. A log likelihood is calculated according to Eq. (3), and the maximum likelihood rule is applied such that the model that generates the highest likelihood is the prediction for which drug condition is associated with heartbeat intervals. We utilize the maximum likelihood rule since for this experiment assume a uniform prior distribution for each drug condition.

## Results

### M. Model Fit

Overlaid KS plots of both models for the number of RR intervals fit, *J*, are shown in Figure 1. We choose *J* = 1500, 3000, and 7000 to be representative of the difference in model fit. The y-axis represents the theoretical percentiles of *F*_*U*_ (·), a uniform distribution on [0, 1], while the x-axis represents the percentiles of *F*_emp_(*τ*_*k*_) from the actual sample data. The red line indicates perfect fit and the dashed lines represent the 95% confidence interval for the data coming from the model. For J = 1500, we can see that the IG model has a very slight advantage in model. However, this advantage disappears as J increases. At J = 7000, we can see that the gamma model outperforms the IG model in goodness of fit, indicating that the gamma model may perform better for larger datasets. While neither model lies within the confidence intervals in the KS plot (indicating that we don’t have a perfect model fit), we note that both model fits are very close to the center diagonal. This indicates to us that both point process models have reasonable fit.

**Fig. 1.**
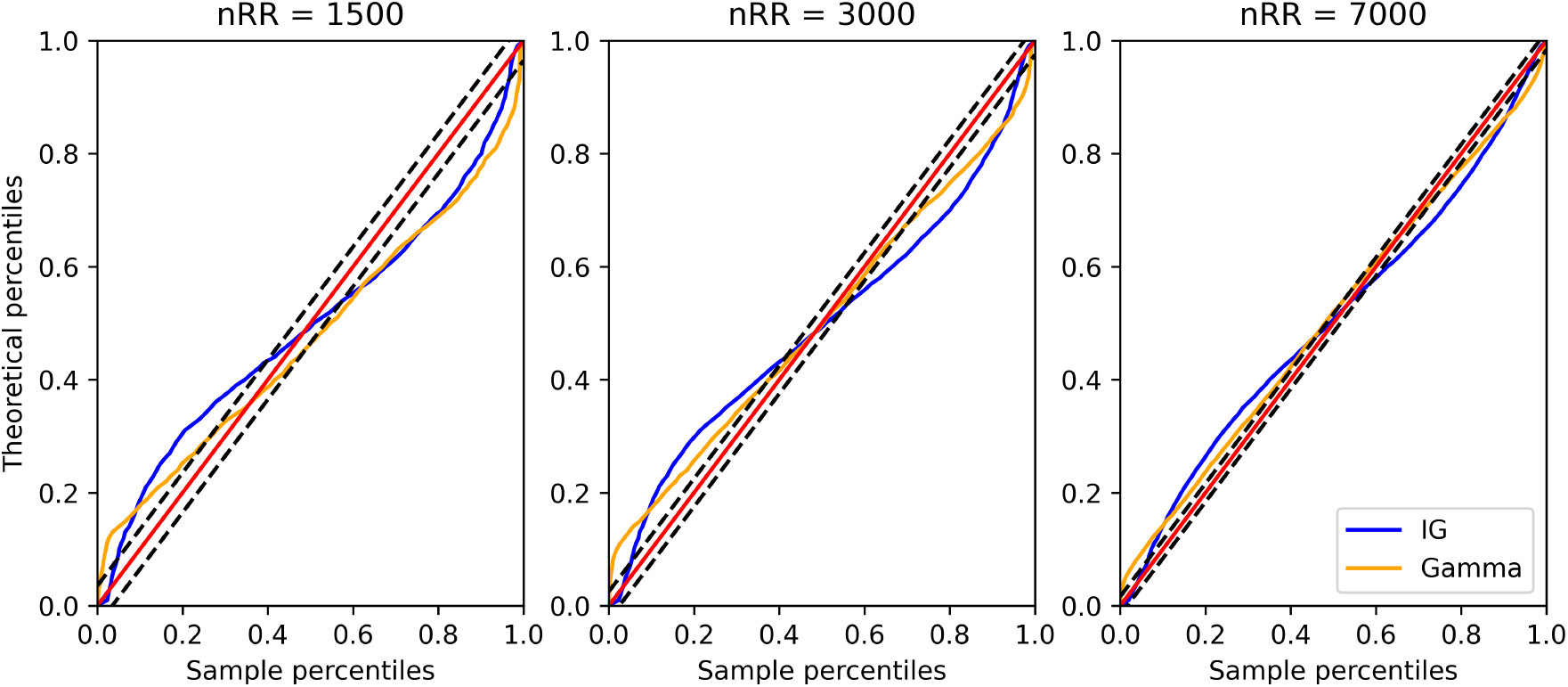
Kolmogorov-Smirnov Plots of time-rescaled ISIs for different numbers of RR intervals fit. In each plot from left to right an increasing number of RR intervals was used to fit both the Inverse-Gaussian and gamma models. The time-rescaling procedure was used which in principle renders its outputs to be unit-rate independent exponentially distributed. The dotted lines are 95% confidence intervals, and the red 45 degree line represents perfect model fit. Notice that as the number of RR intervals fit increases, the gamma model is closer to the center line and achieves better model fit. This suggests that while the gamma model might suffer from worse fit with less data, as the amount of data increases, the gamma model outperforms the Inverse-Gaussian model. From (6)

In Figure 2, the runtime and KS distance for each model are plotted against the number of RR intervals fit. Lower values of KS distance indicate better model fit. In the figure we can see that for all sets of RR intervals fit, the gamma model had a much faster runtime (4x on average). Note, that the gamma model was implemented in Python using CVXPY and the IG model was implemented in MATLAB using the methods described in Barbieri et al (4, 28). For *J >* 1750, the gamma model also has lower KS distances than the IG model, which indicated for larger *J*, the gamma model has better fit than the IG model on this particular dataset.

**Fig. 2.**
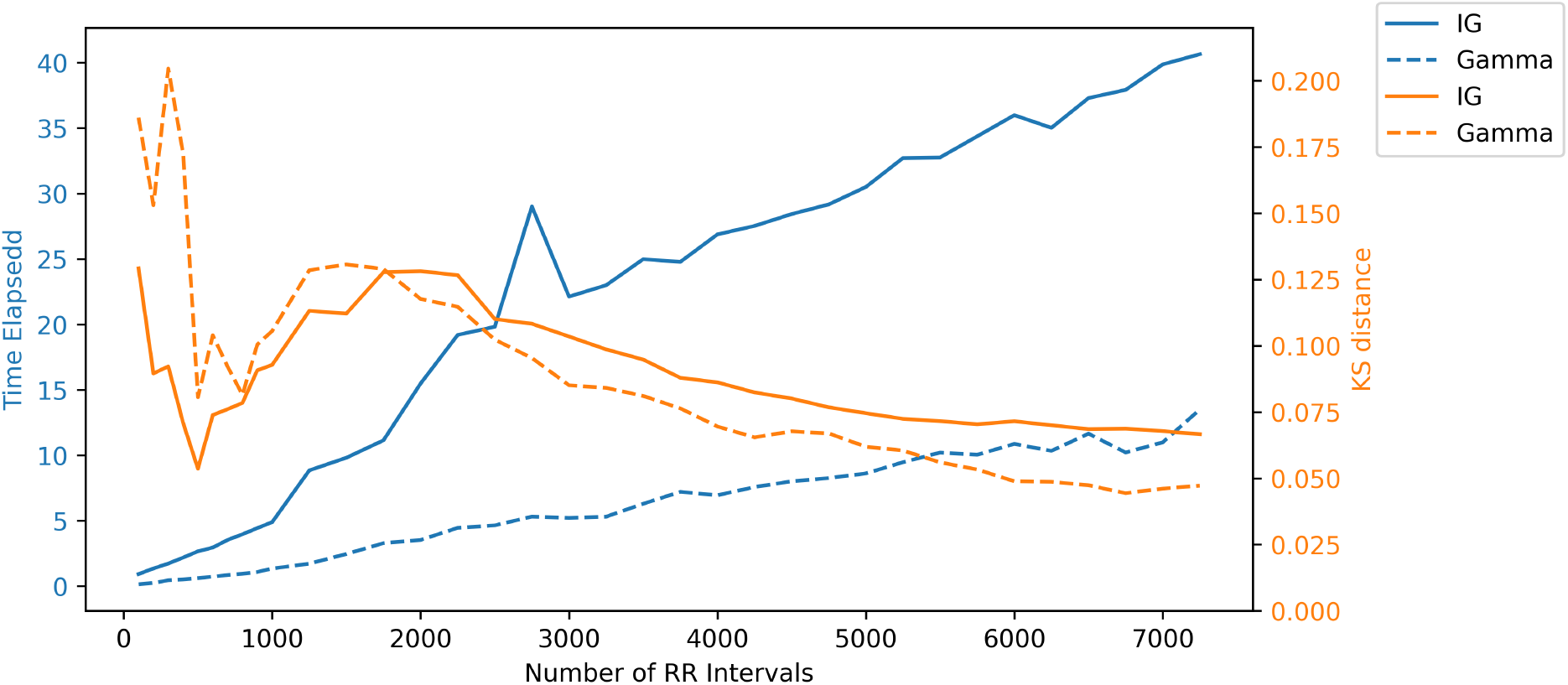
Plot of performance metrics comparing the Inverse-Gaussian to gamma model. The runtime for each algorithm is plotted in blue. For all sets of RR intervals fit, the gamma model ran, on average, 4 times faster. The KS distance (comparing model fit) for each model is plotted in orange. While the gamma model has larger KS distances for small amounts of data, for larger amounts of data the gamma model has a lower KS distance and thus has better model fit. This confirms results in Figure 1. From (6).

### N. Time Series Inference

Results on the time series inference can be found in Fig. 3. Examples for three subjects are shown here for visualization. In each of these plots, the top most trace shows the tilt angle of the tilt table experiment, the middle trace is the heart rate computed from simple 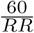 and the bottom trace is the heart rate estimated from the method proposed in this paper. These subjects show a wide variety of responses to the tilt table, where some subjects respond strongly to the tilt (12734, 13960) and one subject does not (12819). By inspection, we note that the mean heart rate estimation computed using our method tracks the fast timevarying dynamics of the heart rate while also being robust to outliers that can be seen by single heart beat timings that deviate from the original trace. Each of these traces also exhibit lower intrinsic variability, better capturing the mean heart rate dynamics. We note that these results are similar to those in Barbieri et al., and we accomplish this while retaining convexity of the problem (4).

**Fig. 3.**
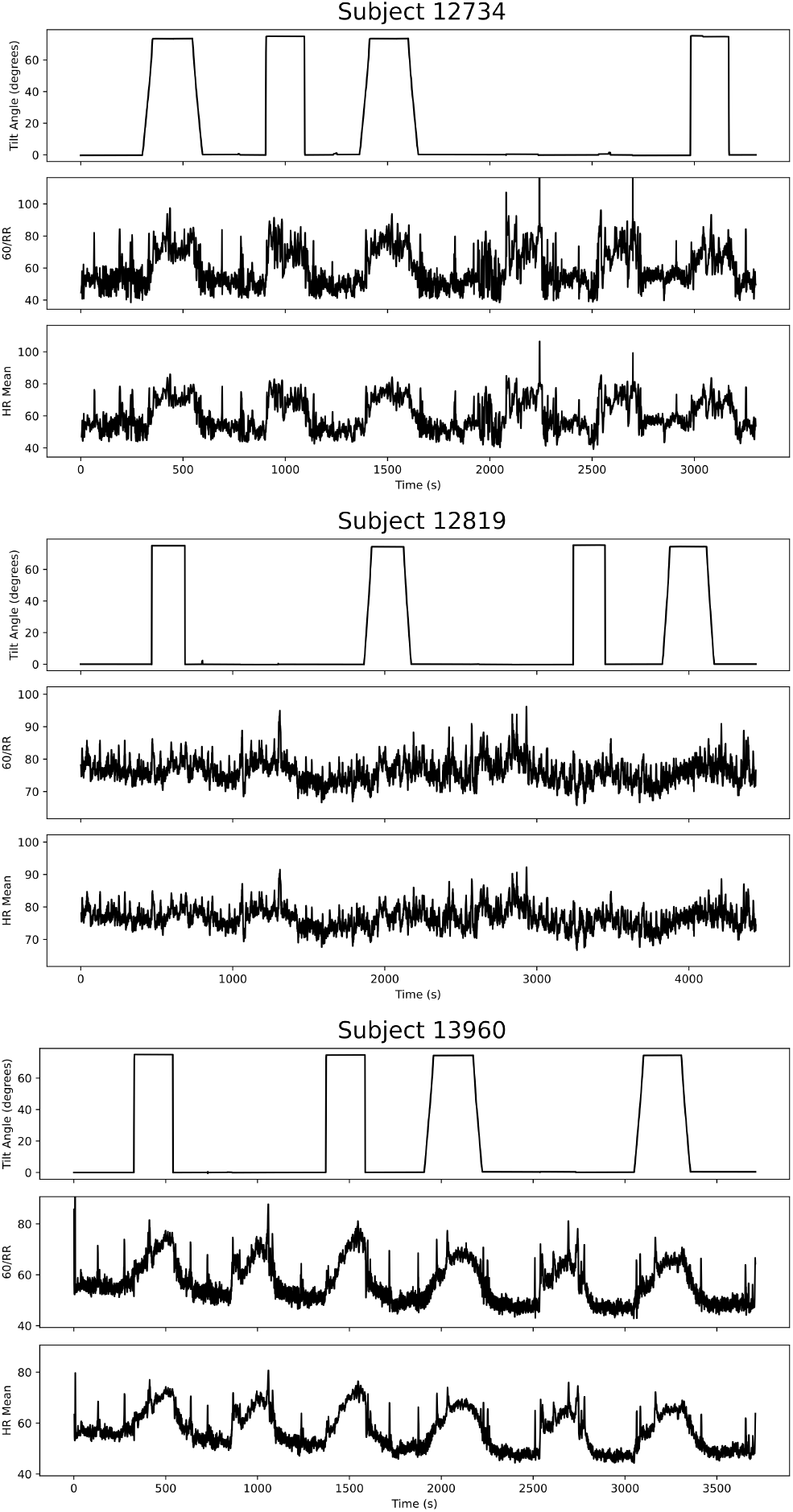
RR Interval Inference. Here we show three selected heart rate time-series estimates from the tilt-table dataset. For each of the three plots, the first trace corresponds to the tilt angle of the subject, the middle trace is 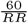, and the last trace is the mean heart rate time series as estimated by the model. Note that the GLM model produces smoother time series estimates and are less influenced by outliers from missed beats or incorrectly identified beats, shown by large spikes away from the trace. The GLM model also does well in tracking the fast transient changes and can operate well in physiologically changing settings.

### O. Sequential Prediction

Fig. 4 shows the results of the sequential prediction algorithm for our methodology and each of the classic machine learning algorithms on the data set for the same three subjects as above. In each quadrant, we plot a scatter plot of the true RR interval against the predicted RR interval. Note, that the variability of the traces above are reflected in the spread of points in these plots. Results immediately show that for all subjects shown in this plot, that out method tracks the best with predicting future heartbeat timings, as the scatter plot consistently lies very close to the 45 degree line that represents perfect prediction. The other classic methods can show good performance on specific instances of data, but scatter plots show that these algorithms often get stuck in local minima and predicted with high bias on the data. For example, observe in subject 12734 how the MLP regressor essentially predicts the same value for an RR interval no matter what the input history or next true RR interval value is.

**Fig. 4.**
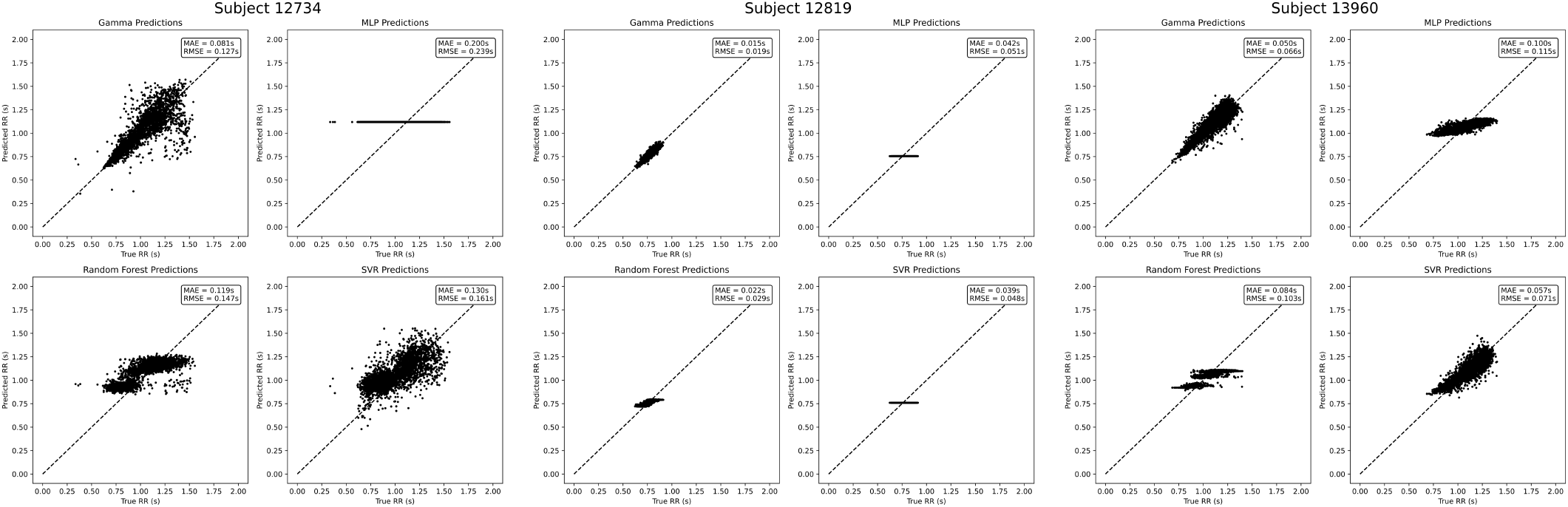
RR Interval Prediction Plots. This figure demonstrates the performance of our GLM method in next beat prediction over the same three example test datasets as in Fig. 3. Each quadrant in each of the three plots represents one of the gamma GLM, MLP, Random Forest, and SVR predictors. The gamma GLM predictor shows high agreement across all subjects with low mean squared error and mean absolute error, even though this dataset exhibits multiple physiological states. Note, that in all of the other methods, the ML predictions and the true next beat timings do not always agree well as they tend to not follow the 45 degree line. This is likely due to the fact that these models either don’t predict well on or get caught in some local optimum.

Also we show in Fig. 5 a boxplot of the model errors for both the MAE and the MSE. In this plot, for each subject the model errors of the each ML model minus the error in the gamma model is plotted. For example, for the MLP column, we include all data points of the form *MSE*_*MLP*_ − *MSE*_*Gamma*_ for the MSE computation. Thus, if the boxplots are positive, we see that on the same dataset, the gamma model has lower error than the ML model and is thus a better predictor. In Fig. 5, we can notice that the MSE and MAE boxplots are almost uniformly above zero, which implies that our model almost always performed better than the standard ML methods in MAE and MSE.

**Fig. 5.**
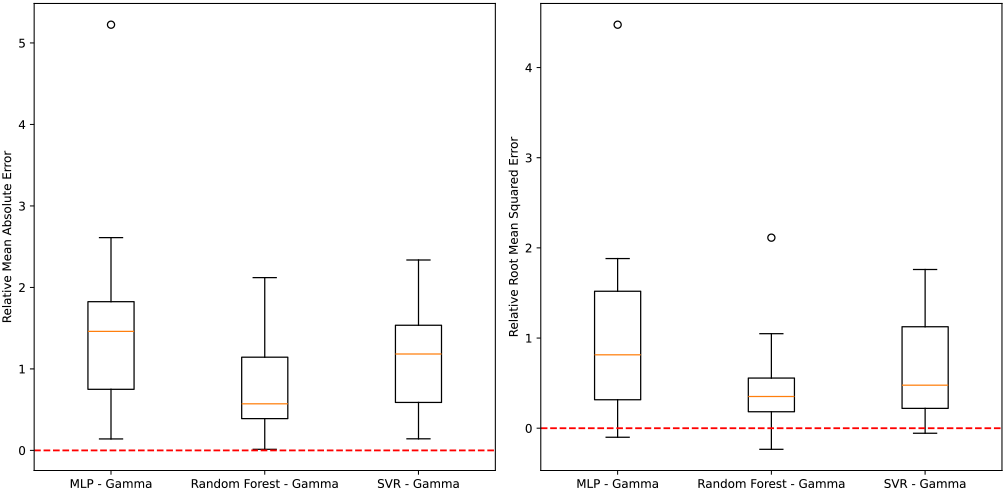
Sequential Prediction Model Errors. The above bar plots show the relative MAE and MSE of each model as compared to the gamma model over all tilt table subjects (n=10). That is each bar is the difference of the error in one of the ML models and the gamma model. Values greater than zero indicate the ML model performs worse than the gamma, and less than zero implies the ML model performs better than the gamma model. Here we see that the ML models uniformly perform worse than the gamma in MAE and nearly so in MSE.

**Fig. 6.**
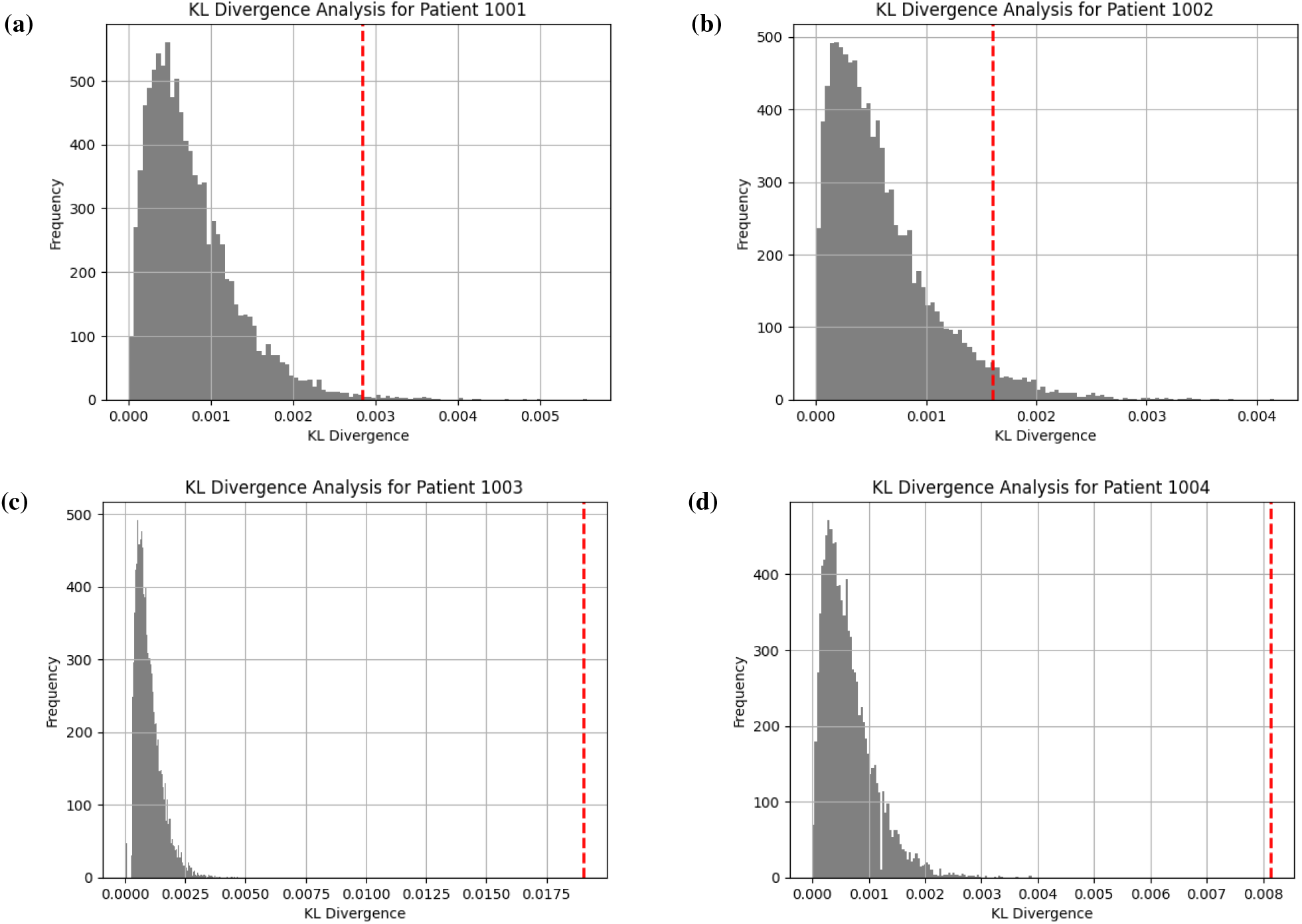
KL divergence plots for Patients: (a) 1001, (b) 1002, (c) 1003, and (d) 1004. Each plot is a histogram of the surrogate KL divergence values generated in surrogate analysis between models fit with drug data and without drug data. The vertical dashed line is the true KL divergence between the models with and without drug data. Each of these histograms are right-skewed distributions. For most patients, the true KL divergence lies far to the right of most of the mass of the distribution, implying that the models fit with and without drug data are significatly different.

### P. Drug Results

The KL divergence bar graphs show a right-skewed distribution with peaks close to 0, indicating that most of the surrogate KL divergence means are small. As expected, this demonstrates that randomly adding regressors does not improve the model performance. However, when correctly encoding drug information, many of the observed KL divergences are nearly an order of magnitude greater than the peak of the surrogate distribution. This difference visually demonstrates that drug data is not just adding noise, but serves as a valuable regressor on an individual basis, improving the predictive power of the model.

Based on the KL divergence analysis experiment, we found that the observed KL divergence means of 19 of the 22 patients were statistically significant compared to the 10000 surrogate KL divergence means generated for each patient. 18 of the 19 statistically significant patients remained significant after performing a Bonferroni correction with the calculated p-values as shown in Table 3. This highlights the robust functionality of the gamma GLM, as it integrates new regressors to better predict heartbeat intervals.

**Table 2.** Observed KL Divergences with Standard Deviation; p-values for Patients were calculated by comparing to surrogate KL divergence means.

| Patient ID | Observed KL Divergences ± std | p-values |
| --- | --- | --- |
| 1001 | 0.0028 ± 0.0022 | 0.0054* |
| 1002 | 0.0016 ± 0.0008 | 0.0506 |
| 1003 | 0.0190 ± 0.0100 | 0.0000* |
| 1004 | 0.0081 ± 0.0088 | 0.0000* |
| 1005 | 0.0011 ± 0.0011 | 0.2383 |
| 1006 | 0.0061 ± 0.0086 | 0.0000* |
| 1007 | 0.0044 ± 0.0038 | 0.0001* |
| 1008 | 0.0218 ± 0.0334 | 0.0000* |
| 1009 | 0.0029 ± 0.0029 | 0.0008* |
| 1010 | 0.0070 ± 0.0091 | 0.0000* |
| 1011 | 0.0419 ± 0.0458 | 0.0000* |
| 1012 | 0.0032 ± 0.0044 | 0.0001* |
| 1013 | 0.0041 ± 0.0049 | 0.0003* |
| 1014 | 0.0066 ± 0.0060 | 0.0000* |
| 1015 | 0.0120 ± 0.0094 | 0.0000* |
| 1016 | 0.0449 ± 0.0538 | 0.0000* |
| 1017 | 0.0246 ± 0.0293 | 0.0000* |
| 1018 | 0.0145 ± 0.0137 | 0.0000* |
| 1019 | 0.0015 ± 0.0012 | 0.1321 |
| 1020 | 0.0089 ± 0.0097 | 0.0000* |
| 1021 | 0.0118 ± 0.0147 | 0.0000* |
| 1022 | 0.0303 ± 0.0340 | 0.0000* |

**Table 3.** Bonferroni-adjusted p-values and significance results.

| Patient ID | Adjusted p-value | ($p_{\text{bon}} < 0.05$)* |
| --- | --- | --- |
| 1001 | 0.1188 | No |
| 1002 | 1.0000 | No |
| 1003 | 0.0000 | Yes |
| 1004 | 0.0000 | Yes |
| 1005 | 1.0000 | No |
| 1006 | 0.0000 | Yes |
| 1007 | 0.0022 | Yes |
| 1008 | 0.0000 | Yes |
| 1009 | 0.0176 | Yes |
| 1010 | 0.0000 | Yes |
| 1011 | 0.0000 | Yes |
| 1012 | 0.0022 | Yes |
| 1013 | 0.0066 | Yes |
| 1014 | 0.0000 | Yes |
| 1015 | 0.0000 | Yes |
| 1016 | 0.0000 | Yes |
| 1017 | 0.0000 | Yes |
| 1018 | 0.0000 | Yes |
| 1019 | 1.0000 | No |
| 1020 | 0.0000 | Yes |
| 1021 | 0.0000 | Yes |
| 1022 | 0.0000 | Yes |

### Q. Predictive Model Results

The drug class predictions were made on single data points consisting of a single heartbeat timing and its history in accordance with the decision rule in Eq. (27) using uniform class priors. The decisions were then compared with the true label of each test time point, and the results are shown in Figure 7. The results show better than chance decisions for all drugs besides Ranolazine, where chance level predictions are 20% accuracy. The accurate predictions occurred best for drugs with longer half-lives (29–32), suggesting that incorporating the loss of drug effects over time would enhance the model’s predictive ability.

**Fig. 7.**
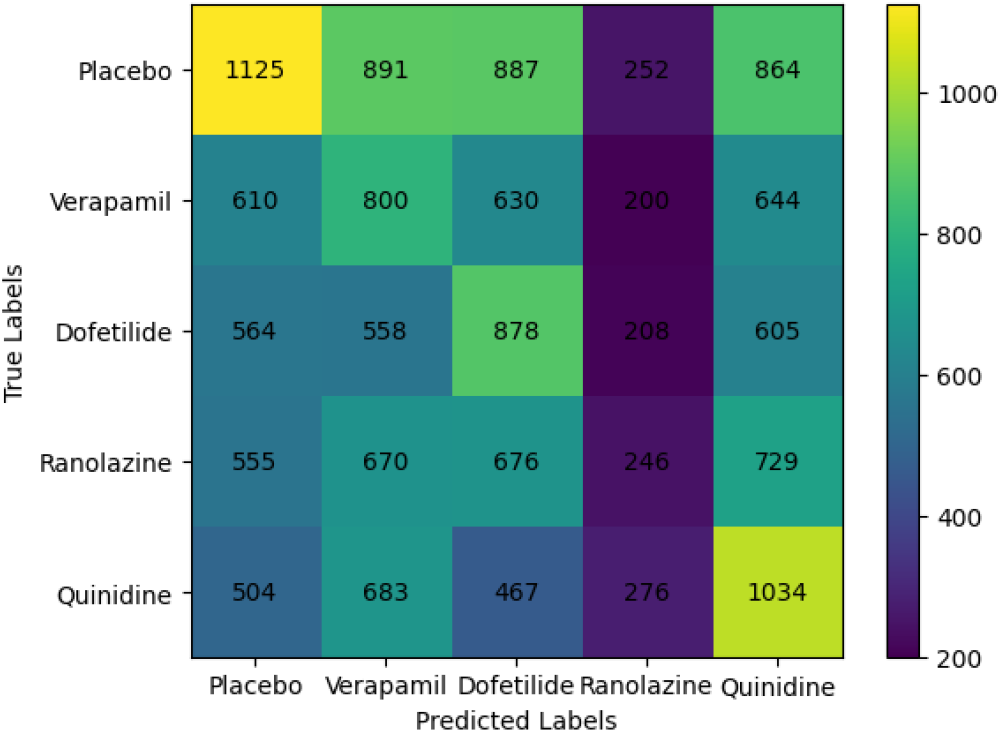
Confusion matrix for drug classification given a single heartbeat timing and history data point. The above figure demonstrates the classification capability of the model by plotting prediction counts of each label, given the true label on the y-axis. Given each row, we demonstrate that for all drugs besides Ranolazine, that the model has higher than chance accuracy in predicting the true drug class given a single heartbeat timing and history. Note, that since we only use a single data point in prediction for this experiment, performance can be significantly enhanced using more data.

## Discussion

In this paper, we have shown that our history-dependent gamma point process model outperforms the original IG point process model in speed and model fit for larger amounts of data. We measured computational speed using runtime of the algorithm, and model fit through the KS distance. Both Figure 1 and Figure 2 validate our model with low runtimes and KS distances. While our original motivation for choosing a gamma GLM to fit the model was in terms of its convexity, for speed and ease of use, we found that it also has the potential for better fit in larger datasets.T further validate this claim, we test on additional datasets.

We also must note that since the gamma distribution was chosen for its convenient computational properties, it may not follow the physiology of the cardiac system as closely as the IG model. This is because the original formulation of the IG model takes into account a Gaussian random walk model of the cardiac membrane potential, which has an IG density describing the interbeat intervals (4). The better model fit of the gamma distribution may in fact be because the gamma distribution is much more flexible in shape than the IG distribution. This means it may be better able to also describe noise or some other unknown physiological process represented in the data, in addition to heartbeat dynamics.

We also demonstrated extensions of our point process model to next heartbeat prediction, information measures, and posterior predictive modeling. In next heartbeat prediction, in comparison to classical machine learning methods, we showed that we were almost uniformly able to predict next heartbeat timings with much lower error. In clinical settings, where subjects were given drugs affecting heart function, we also showed that the CMI was significantly greater than zero, implying that each drug had a differential and detectable effect. We then validated this in the classification setting, where we demonstrated the model’s ability to discriminate which drug a patient had been taking using patient-specific models from EKG data. In this case, we note that we were only using single heartbeat and history pairs to perform the classification and hypothesize that with more data the classifier would be able to better discriminate between drug conditions.

In future work, this approach can be expanded to calculate continuous time estimates of heart rate (HR) and heart rate variability (HRV) as done in the IG model (4)(33). In particular, we envision using state-space methods in order to estimate dynamically changing weights, which will give us an estimate of HR and HRV. For example, using a Kalman filter-like approach would allow for real-time applications of the heartbeat point process model. The key innovation here would be the computational time of the model. The caveat with our model is that for real-time instantaneous measures often times smaller windows of data are required, which is where our model may have worse fit to the data. This implies that while our model may not provide as accurate of an estimate as the IG model in real-time, it would be advantageous for users who desire faster compute times and a simpler algorithm. In addition, in recent literature, a gamma renewal process has shown to arise from a diffusion leaky-integrateand-fire process (34). This indicates to us that in future directions our model may better physiologically describe neural processes and be used for real-time measurements of neural firing rates.

In conclusion, we have developed a convex formulation of the point process heartbeat dynamics model that has lower runtimes and adequate model fit. The development of this model offers an alternative approach to existing methods for easeof-use for non-experts in signal processing to better understand the time-varying dynamics of the cardiovascular system.

## Supplementary Note A: Construction of Drug Regressor matrices

Given that we have a total of 48 disjoint EKGs (triplicates at 16 time points), we need to construct the regressor matrices in a way that doesn’t include, for example, the end of the first EKG as the history of the second EKG. Let *d* = 1, …, 5 be the label of the drug condition and *m* = 1, …, 48 be the EKG recording session. For the model without drug information, we first start by constructing the submatrix for a given drug period and EKG session as

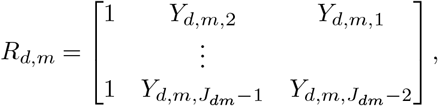

where *J*_*dm*_ represents the total number of beats in just EKG session *m* for drug *d*. In this way, we can keep each recording disjoint. We can then concatenate each of the 48 EKGs and organize by drug such that

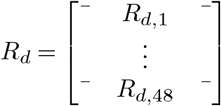

Once we have these regressor matrices grouped by drug, we can generate the final regressor matrix by concatenating those as

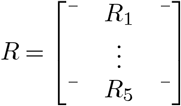

To solve for the parameters, we can then plug into Eq. (30) and Eq. (12) to obtain parameters for the model without explicit drug information denoted by 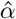 and *ŵ*. For the regressor matrix with the drug information, we can replace the smallest submatrices *R*_*d,m*_ with

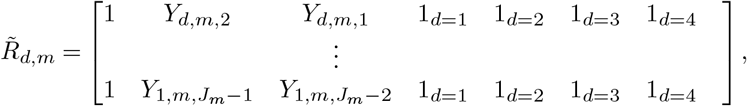

where the drug condition for any specific recording is labeled by a one-hot encoding and the fifth drug condition, *d* = 5, is taken to be all zeros, since it is a placebo condition. For example, if we take data from a the first drug condition (*d* = 1), we can write for any EKG session

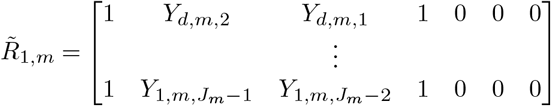

These can be combined for each EKG session in an analogous way to form an 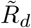 matrix, which can then be combined for each drug condition to form 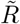.

